# Development of primary osteoarthritis during aging in genetically diverse UM-HET3 mice

**DOI:** 10.1101/2023.12.16.571693

**Authors:** Sher Bahadur Poudel, Ryan R Ruff, Gozde Yildirim, Richard A Miller, David E Harrison, Randy Strong, Thorsten Kirsch, Shoshana Yakar

**Affiliations:** David B. Kriser Dental Center, Department of Molecular Pathobiology, New York University College of Dentistry, New York, NY 10010-4086, USA; David B. Kriser Dental Center, Biostatistics Core, Department of Epidemiology and Health Promotion, New York University College of Dentistry New York, NY 10010-4086; Department of Pathology and Geriatrics Center, University of Michigan, Ann Arbor, MI 48105, USA; The Jackson Laboratory, Bar Harbor, ME 04609, USA; Geriatric Research, Education and Clinical Center and Research Service, South Texas Veterans Health Care System, San Antonio, TX 78229, USA; Barshop Institute for Longevity and Aging Studies and Department of Pharmacology, The University of Texas Health Science Center, San Antonio, TX 78229, USA; Department of Orthopaedic Surgery, NYU Grossman School of Medicine, New York, NY 10100, USA; Department of Biomedical Engineering, NYU Tandon School of Engineering, NY 10010, USA

**Keywords:** osteoarthritis, UM-HET3, sub-chondral bone, cartilage, methylene blue, mitoquinone, antioxidants

## Abstract

This study investigated the prevalence and progression of primary osteoarthritis (OA) in aged UM-HET3 mice. Using the Osteoarthritis Research Society International (OARSI) scoring system, we assessed articular cartilage (AC) integrity in 182 knee joints of 22-25 months old mice.

Aged UM-HET3 mice showed a high prevalence of primary OA in both sexes. Significant positive correlations were found between cumulative AC (cAC) scores and synovitis in both sexes, and osteophyte formation in female mice. Ectopic chondrogenesis did not show significant correlations with cAC scores. Significant direct correlations were found between AC scores and inflammatory markers in chondrocytes, including matrix metalloproteinase-13 (MMP-13), inducible nitric oxide synthase (iNOS), and the NLR family pyrin domain containing-3 (NLRP3) inflammasome in both sexes, indicating a link between OA severity and inflammation. Additionally, markers of cell cycle arrest, such as p16 and β-galactosidase, also correlated with AC scores. Using micro-CT, we examined the correlations between subchondral bone (SCB) morphology traits and AC scores. In male mice, no significant correlations were found between SCB morphology traits and cAC scores, while in female mice, significant correlations were found between cAC scores and tibial SCB plate bone mineral density.

Finally, we explored the effects of methylene blue (MB) and mitoquinone (MitoQ), two agents that affect mitochondrial function, on the prevalence and progression of OA during aging. Notably, MB and MitoQ treatments influenced the disease’s progression in a sex-specific manner. MB treatment significantly reduced cAC scores at the medial knee joint, while MitoQ treatment reduced cAC scores, but these did not reach significance.

In conclusion, our study provides comprehensive insights into the prevalence and progression of primary OA in aged UM-HET3 mice, highlighting the sex-specific effects of MB and MitoQ treatments. The correlations between AC scores and various pathological factors underscore the multifaceted nature of OA and its association with inflammation and subchondral bone changes.

## Introduction

Primary osteoarthritis (OA) refers to the development of OA without any known underlying factors, conditions, or injuries. Aging is a significant risk factor for OA, and during the aging process, there are progressive changes in joint tissues, including articular cartilage, subchondral bone, synovium, and ligaments. These changes involve increased matrix degradation, altered cell metabolism, impaired tissue repair mechanisms [1], and chronic low-grade inflammation [2-5]. In knee OA, which is commonly associated with aging, the articular cartilage gradually deteriorates, becoming thinner and exhibiting structural abnormalities. This compromises its ability to absorb shock and distribute load, resulting in joint pain and dysfunction. The subchondral bone also undergoes remodeling, including thickening and sclerosis, contributing to joint stiffness and further impacting cartilage health. Inflammation in the synovial membrane increases, leading to the production of inflammatory mediators that contribute to cartilage degradation and joint inflammation [6].

Our understanding of primary OA is derived from both clinical and preclinical research. In vivo preclinical animal models that focus mainly on genetic- or injury-induced OA have been particularly useful for two key purposes: investigating the underlying mechanisms of OA and evaluating the effectiveness of various treatment options. However, clinical studies of primary OA have shown variability in the actual onset of the disease, in the rate of OA progression, and in the severity of OA in different individuals. Thus, while inbred mouse models have provided valuable insights into OA mechanisms, they lack the genetic complexity of the human population [7, 8], and provide only limited insights into the natural development of the disease. Studying OA using a single genome compromises generalization and translation of findings [9]. The UM-HET3 mouse model, generated through a specific crossbreeding strategy (**Figure 1A**), offers a genetically heterogeneous population that better represents genetic diversity. The *UM-HET3 mouse model was selected by the Interventional Testing Program (ITP) at the National Institute of Aging (NIA) for studies of anti-aging drugs and was used in several studies to map genes controlling skeletal morphology* [10-13] (https://phenome.jax.org/projects/ITP1). A previous study has shown that UM-HET3 male mice (from one ITP center) developed age-related OA, but exhibited a high degree of variability, showing no significant effects of the tested ITP treatments [14]. In the current study we used 182 stifles of UM-HET3 mice from three ITP centers and explored both sexes.

**Figure 1.**
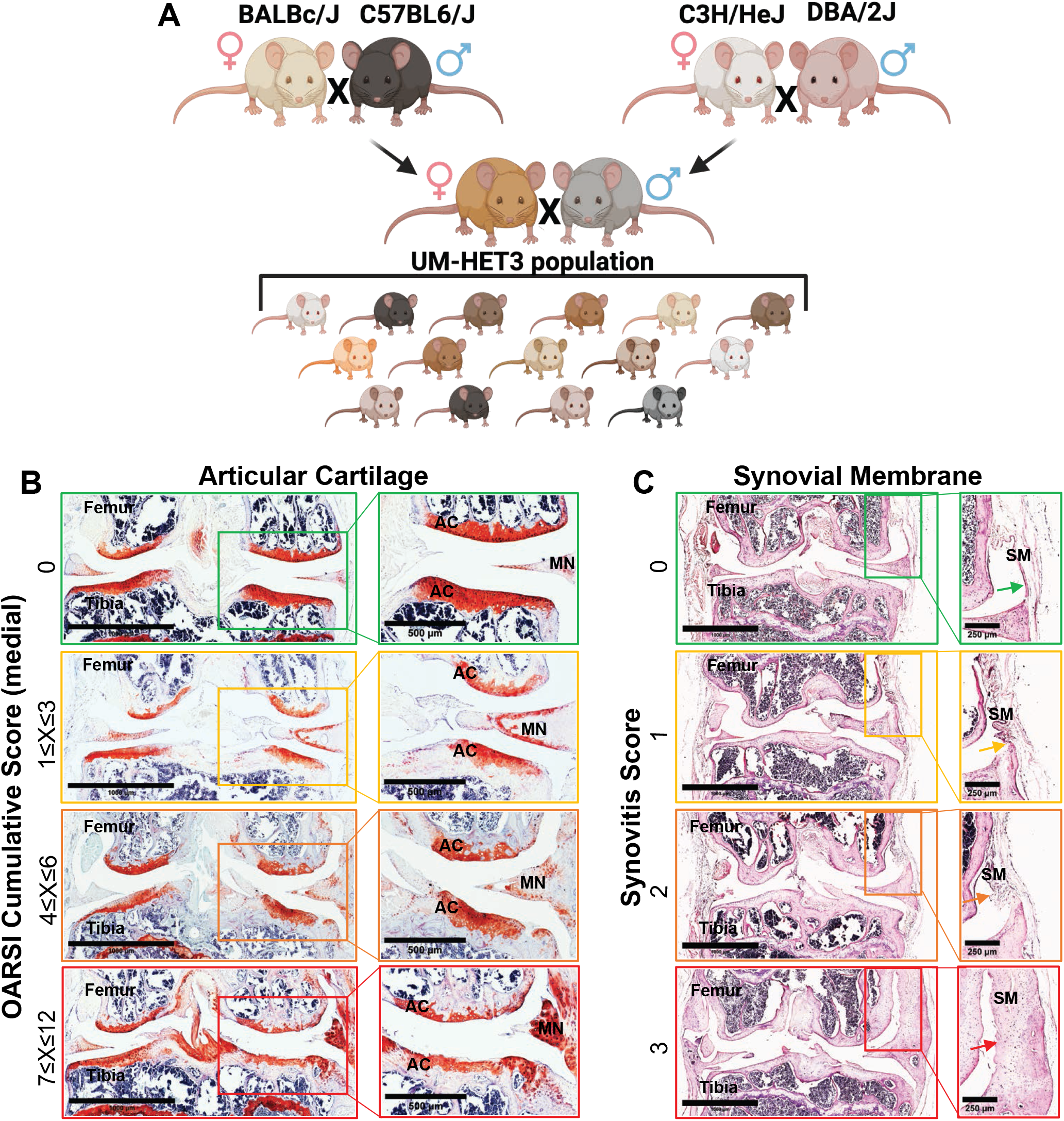
OARSI scoring of stifles from UMHET3 mouse model. (A) UM-HET3 is a genetically diverse mouse model, generated via 4way cross of (BALB/cByJ × C57BL/6J) F1 mothers and (C3H/HeJ × DBA/2J) F1 fathers. Each of the offspring is genetically unique but shares 50% of its genetic material with every other UM-HET3 mouse. (B) Shown are representative knee joint samples that were processed for safranin-O-red, or (C) H&E staining of the synovial membrane that were categorized based on the OARSI scoring system [32-34].

Since OA occurrence intensifies with age, evaluating the presence and severity of OA becomes a crucial factor in assessing the effectiveness of compounds designed to prolong life by influencing pathways that might also play a role in OA development. Therefore, this study aimed to assess the severity of OA associated with aging in genetically diverse male and female mice and to investigate whether the severity of this age-related OA was altered in mice treated with specific agents from the ITP study. Specifically, since common cellular processes are shared between OA and aging, such as mitochondrial dysfunction, oxidative stress, increased inflammation, and accumulation of senescent cells [15], we focused on two compounds that enhance mitochondrial function. Mitochondrial dysfunction and imbalanced production of reactive oxygen species (ROS) are considered contributing factors to OA development [16-18]. Impaired mitochondrial function in chondrocytes is linked to various cellular mechanisms involved in energy regulation, inflammation, and pro-catabolic responses.

Two compounds that affect mitochondrial function were tested by the ITP centers, methylene blue (MB) and mitoquinone (MitoQ). Neither MB nor MitoQ altered lifespan in UM-HET3 mice [19]. MB, an FDA-approved drug used to treat methemoglobinemia, has been shown to enhance mitochondrial function and has demonstrated beneficial effects in animal models of OA [20]. Intra-articular injection of MB in rat and rabbit OA models improved joint symptoms, protected cartilage, and reduced inflammation [21] [22]. MitoQ, a mitochondria-targeted antioxidant [23-25], also suppressed ROS production and ATP synthesis in an osteochondral explant model [26]. Here we report the prevalence of primary OA and the effects of lifelong mitochondrial function enhancement on knee joint tissues during aging in the UM-HET3 mice [27].

## Methods

### Animals

UM-HET3 male and female mice were produced by a cross between (BALB/cByJ × C57BL/6J)F1 mothers (JAX stock #100009) and (C3H/HeJ × DBA/2J)F1 fathers (JAX stock #100004). Detailed housing conditions were specified elsewhere [28]. All experiments were approved by the Institutional Animal Care and Use Committee (IACUC) of each site.

Interventions: MB (28 ppm, starting at 4 months of age) or MitoQ (100 ppm, starting at 7 months of age) were administered in the diet [19]. MB and MitoQ were formulated into irradiated Purina TestDiet 5LG6 diet, and supplied to all three sites such that all sites used the same batches of food.

### Micro computed tomography (micro-CT)

Micro-CT of the knee joints was done in accordance with the American Society for Bone and Mineral Research (ASBMR) guidelines [29]. Intact femur and tibia including the knee joint were scanned using a high-resolution SkyScan micro-CT system (SkyScan 1172, Kontich, Belgium) containing 10-M digital detector set at a 10W energy level (100kV and 100 μA), with a 0.5 mm aluminum filter with a 9.7μm image voxel size. Subchondral bone (SCB) parameters were taken in the distal femur below the AC avoiding cortical bone and included bone volume fraction (bone volume/tissue volume, (BV/TV %), trabecular thickness (Tb.Th, mm), trabecular number (Tb.N, 1/mm), and bone mineral density (BMD, g/cc). Subchondral bone plate (SCBP) thickness and BMD were measured at the proximal tibia. Data reconstruction was done using NRecon software, and data analysis using CTAn software. 3D images were obtained using CT Vox software.

### Histology

Following micro-CT scanning, knee joints were decalcified in 10% EDTA for 3-4 weeks, dehydrated using graded alcohol series and xylene, and processed for paraffin embedding and sectioning. Safranin-O-red staining was used for scoring the AC of the knee joint, including the presence and maturation stage of osteophytes as well as the ectopic chodrogenesis and ossification. H&E staining was used to score inflammation of synovial membrane, and overall morphology.

### Osteoarthritis score

Cartilage damage at medial and lateral tibio-femoral joints was evaluated by two blinded observers using the Osteoarthritis Research Society International (OARSI) scoring system. Loss of cartilage proteoglycan scored by safranin-O-red; normal staining of non-calcified cartilage, scored 0; Loss of safranin-O-red staining without structural changes was scored 0.5; small fibrillations without loss of cartilage was scored 1; Vertical clefts down to the layer immediately below the superficial layer and some loss of surface lamina was scored 2; Vertical clefts/erosion to the calcified cartilage extending to <25% of the articular surface was scored 3; Vertical clefts/erosion to the calcified cartilage extending to 25–50% of the articular surface was scored 4; Vertical clefts/erosion to the calcified cartilage extending to 50–75% of the articular surface was scored 5; Vertical clefts/erosion to the calcified cartilage extending >75% of the articular surface was scored 6. Osteophyte maturity was scored 0 where no osteophytes were detected; 1 when osteophytes were composed of pre-cartilaginous lesion; 2 when osteophytes were composed predominantly of cartilage; 3 when osteophytes were composed of mixed cartilage and bone; and 4 when osteophytes were composed predominantly of bone. Synovitis was scored from H&E-stained sections. Briefly, the thickness of the synovial cell lining layer and the cell density within this layer was scored from 0 to 3, with 0 being the thinnest synovial cell lining layer and low cell density (1-2 cell layers), 1 with 3-5 cell layers, 2 with 6-8 cell layers, and 3 being the thickest synovial cell lining layer (>8 layers) and high cell density. Ectopic chondrogenesis was scored 0 when it was undetectable, 1 when it was found in the synovium and or the capsule, and 2 when it extended into the surrounding ligament and or musculature.

### Immunohistochemistry

Following deparaffinization and rehydration, the tissue sections were treated with citrate-based antigen unmasking solution (H-3300, Vector Laboratories, Inc, California, USA). Then endogenous peroxidase activity was inactivated by BLOXALL endogenous blocking solution (SP-6000, Vector Laboratories, Inc, California, USA). After blocking using the protein block solution (ab64226, Abcam, Massachusetts, USA), the sections were incubated overnight with the primary antibodies at 4°C over-night, stained by rabbit specific HRP/DAB(ABC) detection kit (PK-4001/SK-4100, Vector Laboratories, Inc, California, USA) and counterstained by hematoxylin (Sigma, Missouri, USA)). The images were acquired by DMRXE universal microscope with objective imaging gigapixel montaging workstation (Leica Biosystems, IL, USA) and analyzed by Fiji Image J (Version 1.5r; NIH, Maryland, USA). For IHC, primary antibodies were against iNOS (1:300, PA3030A, Invitrogen, Massachusetts, USA), MMP-13 (1:50, 18165-1-AP, Proteintech, Illinois, USA), NLRP3 (1:300, PA5-88709, Invitrogen, Massachusetts, USA), β-galactosidase (1:400; #ab196838, Abcam), and p16 (1:100; #PA30670, Invitrogen).

### Statistical analysis

Individual descriptive statistics (e.g., means, medians, interquartile ranges) were calculated overall and by group for each outcome. The main effects for treatment and sex, as well as the interaction between the two, were estimated using linear regression (for continuous outcomes) and ordered logistic regression (for categorical outcomes). For each pairwise combination of categorical variables, Spearman’s rank correlation coefficients were computed overall and within treatment and sex, with adjustment for the family wise error rate using the method of Benjamini and Hochberg [30]. Comparisons between sex-specific correlations within each treatment were made using the Fisher r-to-z transformation. Finally, principal component analyses were conducted for continuous multivariate outcomes by treatment and sex, with any instance of missing data being imputed using the regularized iterative PCA algorithm [31]. Data analysis was conducted in R v4.1.3

## Results

### Prevalence of primary OA in UM-HET3 mice

Knee joints of 22-25 months old mice were dissected and processed for safranin-O-red and H&E staining. We used The Osteoarthritis Research Society International (OARSI) scoring system [32-34] to assess the integrity of the AC at the distal femur and proximal tibia (**Figure 1B**) and the state of the synovial membrane at the medial and lateral sides of the joint (**Figure 1C**). The scores of control male and female mice varied both between and within sexes.

Cumulative AC (cAC) scores across *all the knee joint* (*medial and lateral sides* of *both the tibia and the femur*, that can reach a maximal score of 24) revealed significant difference between male and female mice which developed primary OA to varying degrees (OR 2.15, CI [1.289, 3.609], **Figure 2A,B**). However, there was no sex differences within treatment for MB (OR 1.15, CI [0.619, 1.863]) or MitoQ (OR 1.227, CI [0.669, 1.834]).

**Figure 2.**
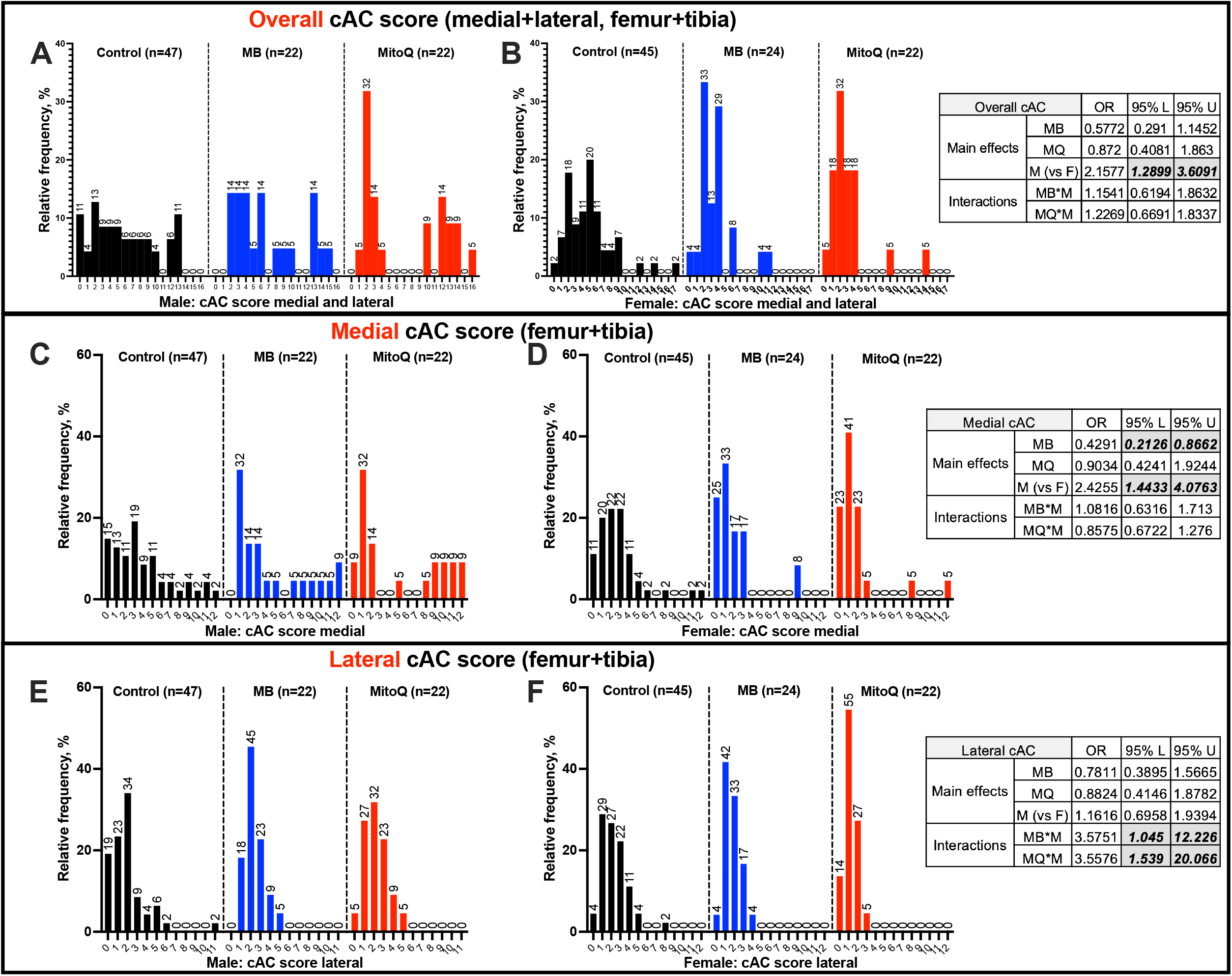
Prevalence and severity of OA in UM-HET3 mice during aging. The frequency of cAC scores at the medial (A,B) and lateral (C,D) sides of the knee joints from control (CTL males n=47, CTL females n=45), MB- (males n=22, females n=24), and MitoQ-treated (males n=22, females n=22) mice. The odd ratio (OR) and the coefficient interval 95% upper (U) and lower (L) of the main effects for treatment and sex, as well as the interaction between the two, were estimated using ordered logistic regression, are indicated in the tables.

cAC scores, specifically at the *medial side* of the tibia and the femur (that can reach a maximal score of 12) revealed significant differences between sexes (OR 2.425, CI [1.443, 4.076], **Figure 2C,D**). Across all mice, treatment with MB significantly reduced the outcomes (OR 0.429, CI [0.213, 0.866]). On the other hand, MitoQ-treated mice did not show significantly different cAC scores at the medial knee joint (OR 0.903, CI [0.424, 1.924]). In females, 11% of control mice showed no OA with age. Treatment with MB or MitoQ increased that ratio to 25% and 23%, respectively, with no effect on the proportion of mice scored 1≤X≤3 in female mice. Approximately 64% of control females, 67% of MB-treated females, and 69% of MitoQ-treated females exhibited an accumulative score of 4≤X≤6 OA.

cAC scores, specifically at the *lateral side* of the tibia and the femur (that can reach a maximal score of 12) revealed no difference between sexes (OR 1.161, CI [0.695, 1.939], **Figure 2E,F**). Approximately 66% of control males, 86% of MB-treated males, and 82% of MitoQ-treated males exhibited a cAC score of 1≤X≤3 OA in the lateral side. In females, treatment with MitoQ associated with 14% of cAC score of 0 and 87% of cumulative AC score of 1≤X≤3. Overall, we found that at the lateral side of the knee joint, treatment significantly differed between sexes. Male mice had 3.575X the odds of female mice of having a high cAC scores (OR 3.575, CI [1.045, 12.226]). Likewise, MitoQ-treated male mice had 3.557X higher odds (OR 3.557, CI [1.539, 20.066]) of having a high cAC score than females

### AC scores correlate with osteophytosis, synovitis and ectopic chondrogenesis

We investigated the relationships between cAC scores and three other factors: synovitis (**Figure 3A,B**), osteophytosis (**Figure 3C,D**) and ectopic chondrogenesis (Ec.Chond) (**Figure 3E,F**). There was no significant difference in the correlation between AC score and synovitis in males versus females (**Figure 3A,B**). In the MitoQ group, but not MB, there were significant differences (p=0.0117) in the correlations between cAC and synovitis in males compared to females (**Figure 3G)**.

**Figure 3.**
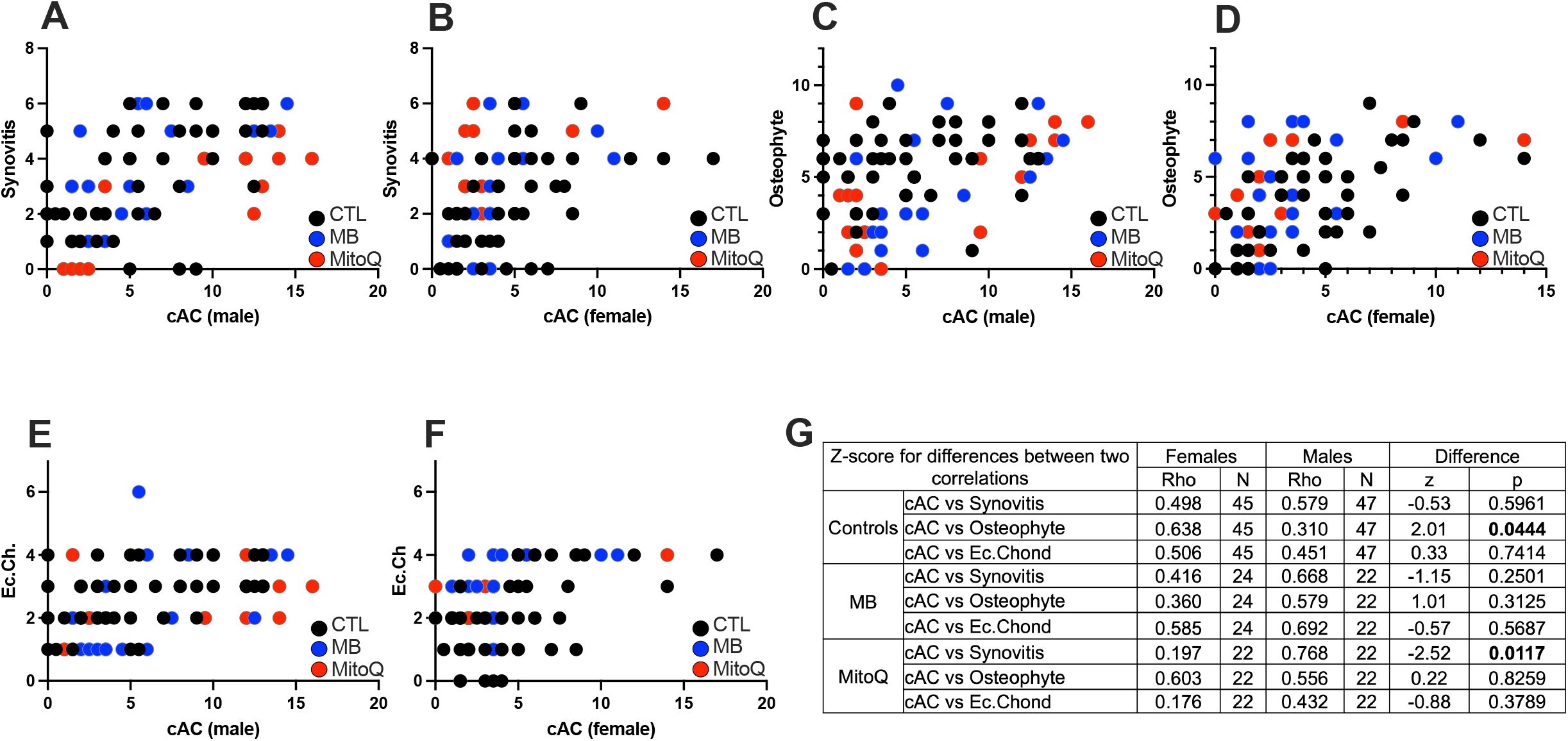
Correlations between cumulative AC (cAC) scores and synovitis, osteophytosis, or ectopic chondrogenesis (Ec.Chond). Correlations between cAC scores and synovitis in male (A) and female (B) mice. Correlations between cAC scores and osteophyte formation in male (C) and female (D) mice. Correlations between cAC scores and ectopic chondrogenesis in in male (E) and female (F) mice. A summary table of the Z score differences between each correlation along with the p values (G). CTL males n=47, CTL females n=45, MB males n=22, MB females n=24, MitoQ males n=22, MitoQ females n=22.

We used a semiquantitative method to score the degree of osteophytes present in the medial and lateral sides of the knee joint. In male mice, osteophytosis did not correlate with cAC scores (**Figure 3C,D**). However, there were significant differences in the correlations between AC score and ostephytosis in males as compared to females (p=0.044) (**Figure 3G)**. Lastly, there were no correlations between ectopic chondrogenesis and cAC scores in either sex for mice treated with MitoQ or MB (**Figure 3E,F**).

### AC scores correlate with markers of inflammation in chondrocyte

We used immunohistochemistry (IHC) approach to determine the levels of inflammatory markers in chondrocytes of the AC. Three inflammatory markers were selected including MMP-13, iNOS, and the NLRP3 inflammasome. We found significant direct correlations between the AC scores and the protein levels of MMP-13, iNOS, and NLRP3 in chondrocytes at the medial part of the knee joint in both male and female mice (**Figure 4A, supplement figure 1**). Similar correlations were found in the lateral part of the knee joint, although, not all were significant (**Figure 4B**). As expected, synovitis score correlated with the protein levels of iNOS, and NLRP3 at the synovial membrane at the medial part of the joint (**Figure 4C, supplement figure 2**). Finally, we found significant direct correlations between AC scores and the protein levels of markers of cell cycle arrest including p16 and β-galactosidase in chondrocytes (**supplement figure 3**).

**Figure 4.**
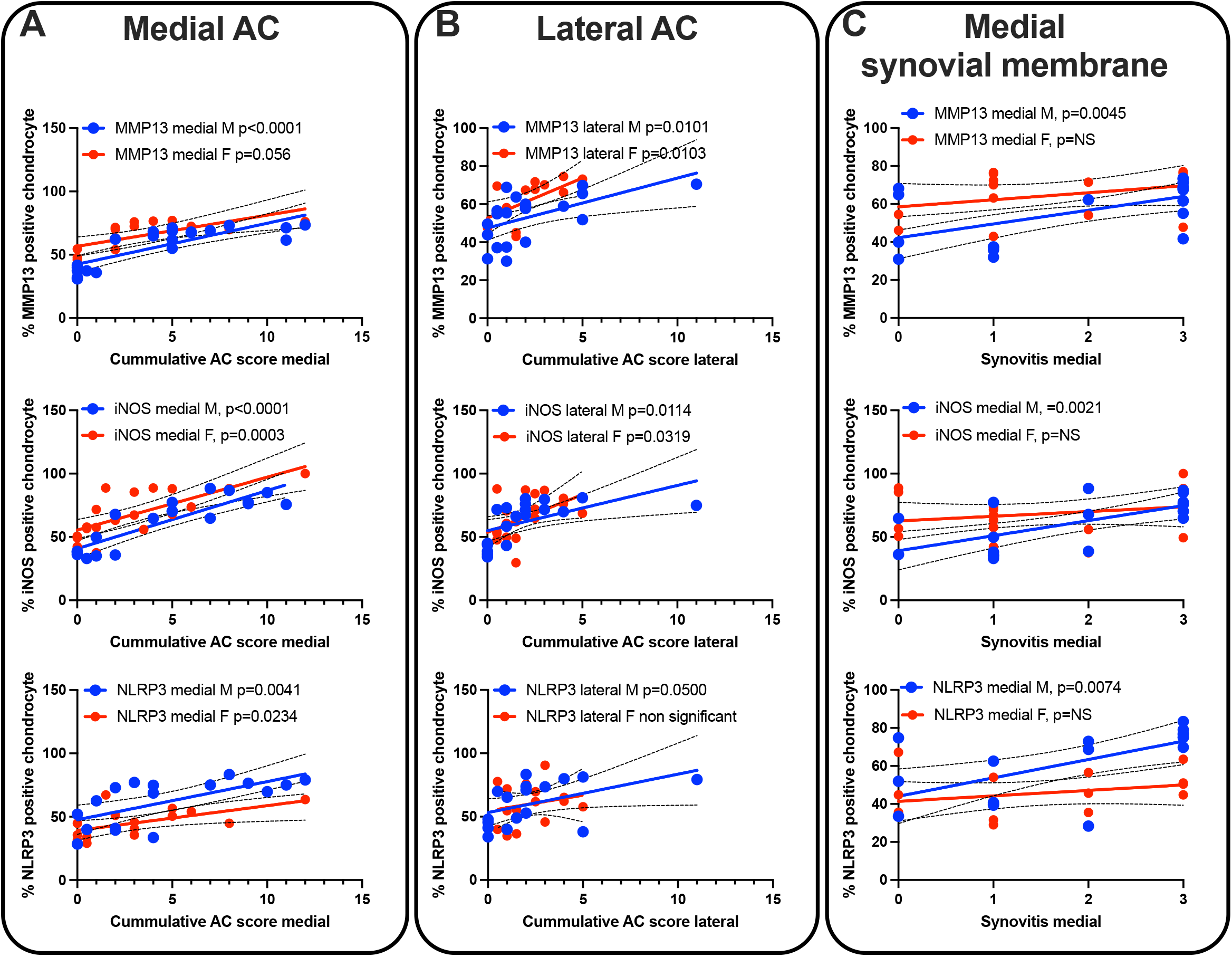
Correlations between cumulative AC scores and markers of inflammation. Correlations between cumulative AC scores at the medial (A) and lateral (B) side of the joint with the levels of MMP13 (males n=21, females n=15), iNOS (males n=19, females n=19), and NLRP3 (males n=17, females n=15) in AC chondrocytes. (C) Sections of the medial synovial membrane in control mice immunostained with iNOS (males n=19, females n=19), and NLRP3 (males n=17, females n=15).

### SCB morphology in aged UM-HET3 mice was not modified by MB or MitoQ treatment

We examined the morphology of the subchondral bone (SCB) and subchondral plate (SCBP) using micro-CT analysis (**Table 1, supplement figure 4**). All parameters tested, including SCB volume (bone volume/total volume, BV/TV) in femur (**Figure 5A**) or tibia (**Figure 5E**), SCB trabecular thickness (Tb.Th) in femur (**Figure 5B**) or tibia (**Figure 5F**), or subchondral bone mineral density (BMD) in femur (**Figure 5C**) or tibia (**Figure 5G**) significantly deferred between sexes but were not altered with treatment. Similarly, tibial SCBP Th (**Figure 5H**) or BMD (**Figure 5I**), significantly deferred between sexes, and MitoQ treatment significantly increased SCBP BMD (p=0.0438) compared to controls. We found that morphological traits of the SCB in the femur correlated significantly to those of the tibia, indicating that both SCB compartments are regulated in a similar manner (**supplement figure 5**).

**Table 1.** SCB morphology of male and female mice determined by micro-CT. Bone volume/total volume (BV/TV), trabecular thickness (Tb.Th), and bone mineral density (BMD).

|  |  |  |  |  |  |  |  | Main effects |  |  | Interactions |  |
| --- | --- | --- | --- | --- | --- | --- | --- | --- | --- | --- | --- | --- |
|  | M CTL<br>(n=47) | M MB<br>(n=22) | M MitoQ<br>(n=22) | F CTL<br>(n=45) | F MB<br>(n=24) | F MitoQ<br>(n=22) |  | MB | MQ | M (vs F) | MB*M | MQ*M |
| Distal Femur Subchondral Bone |  |  |  |  |  |  |  |  |  |  |  |  |
| BV/TV, % | 21.7618±<br>0.7489 | 20.9130±<br>1.0306 | 21.8811±<br>1.5200 | 17.6319±<br>0.6837 | 17.5472±<br>0.7915 | 17.7906±<br>0.9468 | B | -0.526 | 0.2029 | 3.924 | -0.76258 | -0.03787 |
|  |  |  |  |  |  |  | SE | 1.043 | 1.0497 | 0.7396 | 1.79945 | 1.85574 |
|  |  |  |  |  |  |  | p | 0.615 | 0.847 | 3.39E-07 | 0.67225 | 0.983741 |
| Tb.Th,<br>mm | 0.0688±<br>0.0012 | 0.0678±<br>0.0020 | 0.0687±<br>0.0020 | 0.0749±<br>0.0016 | 0.0759±<br>0.0022 | 0.0789±<br>0.0023 | B | -0.000145 | 0.0019519 | -0.0075682 | -0.0019332 | -0.0041596 |
|  |  |  |  |  |  |  | SE | 0.00207 | 0.0020854 | 0.0014693 | 0.0035634 | 0.0036748 |
|  |  |  |  |  |  |  | p | 0.944 | 0.351 | 6.97E-07 | 0.58817 | 0.25924 |
| Tb.Sp,<br>mm | 0.2532±<br>0.0057 | 0.2539±<br>0.0065 | 0.2512±<br>0.0109 | 0.3296±<br>0.0065 | 0.3243±<br>0.0061 | 0.3336±<br>0.0096 | B | -0.0022881 | 0.0007131 | -0.0762672 | 0.0060545 | -0.0059524 |
|  |  |  |  |  |  |  | SE | 0.0084013 | 0.0084558 | 0.0059576 | 0.0144828 | 0.0149359 |
|  |  |  |  |  |  |  | p | 0.786 | 0.933 | 2.00E-16 | 0.676 | 0.691 |
| Tb.N,<br>1/mm | 3.1403±<br>0.0790 | 3.0609±<br>0.0990 | 3.1812±<br>0.1763 | 2.3486±<br>0.0742 | 2.3010±<br>0.0653 | 2.2506±<br>0.1004 | B | -0.062927 | -0.024334 | 0.816088 | -0.03187 | 0.138898 |
|  |  |  |  |  |  |  | SE | 0.110956 | 0.111676 | 0.078683 | 0.191178 | 0.197159 |
|  |  |  |  |  |  |  | p | 0.571 | 0.828 | <2e-16 | 0.868 | 0.482 |
| BMD,<br>g/cc | 0.2480±<br>0.0072 | 0.2344±<br>0.0090 | 0.2530±<br>0.0137 | 0.2208±<br>0.0072 | 0.2114±<br>0.0087 | 0.2307±<br>0.0097 | B | -0.00532 | 0.001547 | 0.025015 | -0.004436 | -0.00509 |
|  |  |  |  |  |  |  | SE | 0.010117 | 0.010183 | 0.007175 | 0.017461 | 0.018007 |
|  |  |  |  |  |  |  | p | 0.599648 | 0.879432 | 0.000619 | 0.79975 | 0.77775 |
| Proximal Tibia Subchondral Bone |  |  |  |  |  |  |  |  |  |  |  |  |
| BV/TV, % | 20.3482±<br>0.6811 | 20.6231±<br>1.2275 | 18.6409±<br>1.0443 | 17.0072±<br>0.7781 | 16.7714±<br>1.0041 | 15.9309±<br>0.7885 | B | -1.1174 | -0.4082 | 3.3361 | 0.46492 | -0.67686 |
|  |  |  |  |  |  |  | SE | 1.0344 | 1.0038 | 0.7198 | 1.76073 | 1.77353 |
|  |  |  |  |  |  |  | p | 0.2815 | 0.6848 | 7.09E-06 | 0.79207 | 0.70321 |
| Tb.Th,<br>mm | 0.0651±<br>0.0010 | 0.0671±<br>0.0015 | 0.0642±<br>0.0013 | 0.0724±<br>0.0014 | 0.0705±<br>0.0018 | 0.0709±<br>0.0018 | B | -0.001612 | 0.0002615 | -0.006148 | 3.83E-03 | 5.71E-04 |
|  |  |  |  |  |  |  | SE | 0.0016909 | 0.001641 | 0.0011768 | 2.87E-03 | 2.89E-03 |
|  |  |  |  |  |  |  | p | 0.3418 | 0.8736 | 5.06E-07 | 0.1828 | 0.8434 |
| Tb.Sp,<br>mm | 0.2135±<br>0.0030 | 0.2180±<br>0.0063 | 0.2314±<br>0.0057 | 0.2601±<br>0.0038 | 0.2558±<br>0.0063 | 0.2674±<br>0.0056 | B | 0.00244 | 0.010634 | -0.041754 | 0.008938 | 0.010821 |
|  |  |  |  |  |  |  | SE | 0.005548 | 0.005384 | 0.003861 | 0.009407 | 0.009475 |
|  |  |  |  |  |  |  | p | 0.6607 | 0.0499 | <2e-16 | 0.343 | 0.255 |
| Tb.N,<br>1/mm | 3.1047±<br>0.0739 | 3.0392±<br>0.1355 | 2.8963±<br>0.1369 | 2.3249±<br>0.0827 | 2.3618±<br>0.1231 | 2.2370±<br>0.0891 | B | -0.10127 | -0.07336 | 0.72489 | -0.10591 | -0.12402 |
|  |  |  |  |  |  |  | SE | 0.11812 | 0.11463 | 0.0822 | 0.20097 | 0.20244 |
|  |  |  |  |  |  |  | p | 0.392 | 0.523 | 1.34E-15 | 0.599 | 0.541 |
| BMD,<br>g/cc | 0.2389±<br>0.0070 | 0.2365±<br>0.0105 | 0.2302±<br>0.0103 | 0.2175±<br>0.0075 | 0.2080±<br>0.0105 | 0.2085±<br>0.0082 | B | -0.010901 | -0.004595 | 0.0234 | 6.80E-03 | 6.80E-03 |
|  |  |  |  |  |  |  | SE | 0.01027 | 0.009966 | 0.007147 | 1.75E-02 | 1.76E-02 |
|  |  |  |  |  |  |  | p | 0.28999 | 0.64532 | 0.00128 | 0.698 | 0.9957 |
| Tibia Subchondral Plate |  |  |  |  |  |  |  |  |  |  |  |  |
| SCB plate<br>Th, mm | 0.1080±<br>0.0036 | 0.0918±<br>0.0055 | 0.1027±<br>0.0044 | 0.1009±<br>0.0032 | 0.0890±<br>0.0061 | 0.1059±<br>0.0044 | B | -0.007202 | -0.005887 | 0.003244 | -0.0037615 | -0.0097749 |
|  |  |  |  |  |  |  | SE | 0.004989 | 0.004816 | 0.003462 | 0.0084504 | 0.008512 |
|  |  |  |  |  |  |  | p | 0.1507 | 0.2232 | 0.3501 | 0.6568 | 0.2524 |
| SCB plate<br>BMD,<br>g/cc | 0.8638±<br>0.0147 | 0.8529±<br>0.0168 | 0.9381±<br>0.0058 | 0.8914±<br>0.0167 | 0.8690±<br>0.0190 | 0.9620±<br>0.0085 | B | 0.02725 | 0.03507 | -0.02518 | 0.01436 | 0.00654 |
|  |  |  |  |  |  |  | SE | 0.01429 | 0.01789 | 0.01727 | 0.0304 | 0.03062 |
|  |  |  |  |  |  |  | p | 0.1295 | 0.0438 | 0.0441 | 0.6374 | 0.8312 |

**Figure 5.**
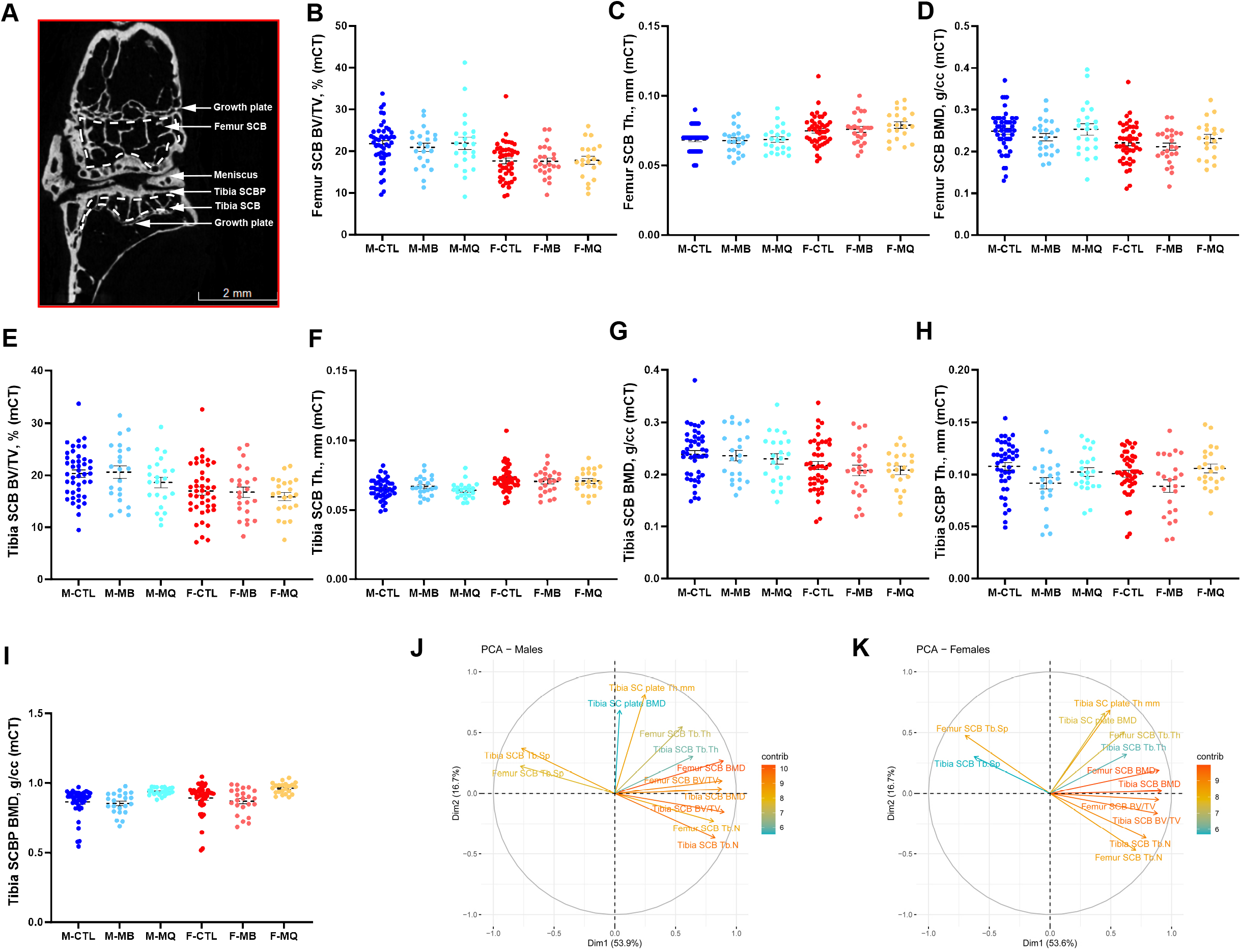
Subchondral bone (SCB) and subchondral plate (SCP) morphology analyzed by micro-CT. (A) 2D image of a micro-CT scan indicating the knee joint regions of interest that were analyzed including the SCB of the distal femur, SCB of the proximal tibia, and the SCB plate of the proximal tibia. Parameters assessed by micro-CT included femur (B) bone volume/total volume (BV/TV), (C) trabecular thickness (Tb.Th), and (D) SCB mineral density (BMD). Likewise, tibia BV/TV (E), Tb.Th of the tibia SCB (F), and tibia SCB mineral density (G). SCB plate (SCBp) thickness (H), and SCBp bone mineral density (I), were determined at the proximal tibia. Data presented as mean ± SEM, p values are indicated. CTL males n=44, CTL females n=44, MB males n=22, MB females n=24, MitoQ males n=22, MitoQ females n=22. SCB phenotypic measurements from PCA of both male (J) and female (K) mice. Positively correlated variables point to the same side of the plot; negatively correlated variables point to opposite sides of the plot. Length and color of arrows indicate contribution of the variable to the overall variance of the data. Arrows that are close together are strongly positively correlated, arrows that are about 90 degrees (orthogonal) to each other are uncorrelated, and those far away (such as 180 degrees) are strongly negatively correlated.

Results from principal component analyses (PCA) suggest that much of the variation in micro-CT measures across samples occurs in two dimensions (70%) (**Figure 5J,K)**. In both males and females, the strongest influencing measurements included femur SCB BV/TV, femur SCB BMD, and tibia SCB BV/TV for the first component, while femur SCB Tb.Th had substantial influence on the second component. There was a strong positive correlation between femur and tibial measurements with the exception of strong negative correlations with both tibia and femur SCB Tb.Sp. Surprisingly, PCA indicate that measurements for the tibia SCBP were largely orthogonal. In males, contributions from tibia SCB Tb.Th and tibia SCBP BMD were not as strong, nor were tibia SCB Tb.Th and tibia SCB Tb.Sp in females. Finally, we did not find significant interactions between SCBP morphology and the cAC scores at the lateral or medial of the tibia (**supplement figure 6**).

## Discussion

OA, a widespread and chronic joint disorder, is marked by ongoing deterioration of cartilage, alterations in subchondral bone, bone marrow lesions, damage to the meniscus, and synovitis. In the current study we utilized 182 stifles of genetically diverse UM-HET3 mice to model the prevalence and severity of primary OA during aging in male and female mice. We found that at the medial side of the knee joint 90% of control female and 85% of control male mice developed primary OA to varying degrees, accompanied by synovitis, osteophytosis, and calcified menisci. Interestingly, 64% of control females showed cAC scores of 1≤X≤3, and only 43% of males showed scores at that range in the medial side of the knee joint.

Both male and female control mice showed significant correlations between AC scores and synovitis. In control female mice we found significant correlation between osteophytosis and AC scores. Notably, osteophytes contribute to both the functional properties of affected joints and clinically relevant symptoms. They are closely associated with cartilage damage, although they can also develop without explicit cartilage damage [35]. The precursor cells for osteophyte formation are debatable and include mesenchymal stem cells in the periosteum [36] or synovium-derived cells [37]. The TGFβ superfamily of growth factors and macrophages play important roles in osteophyte induction [38]. In control males, osteophytosis did not correlate with AC scores.

The progression of osteoarthritis, triggers inflammatory reaction, characterized by increased expression of the inflammatory markers in AC chondrocytes and cells of the synovial membrane. Specifically, we report significant correlations between protein levels of MMP13, iNOS, and NLRP3 in AC chondrocytes and synovial membrane at the medial knee joint in both male and female mice. Release of inflammatory factors interfere with the anabolic and catabolic activities of chondrocytes and may lead to cell death [39, 40].

Previous studies have reported that mitochondrial dysfunction and oxidative stress contribute to OA development [41]. In chondrocytes, the production of mitochondrial ROS has been linked to a marked increase in cyclooxygenase, a powerful catabolic inflammatory agent [42] and MMP-13 [43], highlighting their potential role in degradation of the AC. Earlier research has established the significant role of oxidants in modulating metabolic processes in cartilage [44-46]. Together, it is expected that inhibiting excessive ROS accumulation can protect against chondrocyte damages and slow down OA progression. Further, mitochondria in chondrocytes are mechanically connected to the cell membrane through f-actin, facilitating their movement and distribution in the cytosol [47]. This connection between mitochondria and the cytoskeleton plays a crucial role in managing energy and ROS production [45], as well as maintaining calcium balance [48] and controlling mitochondrial apoptotic factors [49]. In light of these evidence, we studied the effects of lifelong treatment with MB or MitoQ on the natural history of primary OA in UM-HET3 mice.

MB and MitoQ affect oxidative stress in tissues of the knee joint via different mechanisms. MB ameliorates oxidative stress via stimulation of Nrf2 and its translocation to the nucleus in both in chondrocytes and synoviocytes and exerts anti-inflammatory effects in the synovium and neurons that innervate the knee joint [21]. Evidence from the current study indicate that lifelong treatment with MB lowered the cAC scores compared to controls, suggesting that it may alter the natural history of primary OA in UM-HET3 mice. MitoQ, on the other hand, functions as a mitochondrial antioxidant [50]. Its lipophilic moiety allows MitoQ to concentrate within mitochondria up to a thousand times more than other general antioxidants [51]. MitoQ is known for its protective effects against diseases related to oxidative damage. In a study using a model of bovine cartilage explants subjected to mechanical stress, MitoQ significantly reduced the production of ROS [26]. Furthermore, MitoQ treatment was found to mitigate the breakdown of the extracellular matrix (ECM), reduce oxidative stress, and lessen the inflammatory response in chondrocytes. This was observed in a mouse model of OA induced by destabilizing the medial meniscus (DMM) [52] and was likely due to its activation of the NRF2 pathway and the promotion of mitophagy in chondrocytes. In the current study we found that MitoQ-treated male mice had higher lateral, but not medial, cAC scores compared to female mice. Additionally, mice treated with MitoQ, regardless of sex, showed lower measures for SCB Plate BMB in Tibia.

In this study, we report on the prevalence and severity of primary OA in the genetically diverse UM-HET3 mice. We also examined how treating these mice with the antioxidants MB and MitoQ from early adulthood into old age impacts the development of primary OA. Our research has certain limitations, including the fact that we only examined mice at one age range (22-25 months). We established the validity of our model by showing correlations between OA features and indicators of inflammation, synovitis, and osteophyte formation. However, our study did not explore the direct causes of these conditions. We also recognize that our research does not delve into the cellular or molecular mechanisms that might trigger or advance OA. Nevertheless, our findings demonstrate that primary OA in UM-HET3 mice reflects the prevalence and severity seen in human primary OA. This suggests that these mice are a suitable model for testing systemic treatments that could influence aging processes, particularly in the context of developing primary OA.

## Author contribution

Conceptualization: SY

Funding acquisition: SY

Formal analyses: RRR, SY

Investigation: SBP, GY

Statistical analyses: RRR

Resources: RAM, DEH, RS

Writing review and editing: SY, TK

## Acknowledgements

Financial support received from the National Institutes of Health Grant R01AG056397 and B01 (2024) Department of Molecular Pathobiology Accelerator Award to SY. SBP is supported by New York University Provost’s Postdoctoral Fellowship Program. This work was also supported by the National Institutes of Health grant to The Jackson Laboratory Nathan Shock Center of Excellence in the Basic Biology of Aging AG038070, U01-AG022303 to RAM, UO1-AG022308 to DEH, U01-AG013319 to RLS, RLS is supported by a Senior Research Career Scientist Award from the Department of Veterans Affairs Office of Research and Development, and S10 OD010751-01A1 for micro-computed tomography.

## Co-authors details

## Supplement Figure Legends

**Supplement Figure 1:** Shown are representative knee joint sections stained with MMP-13, iNOS, and NLRP3 antibodies in both male and female mice. Quantification is available in figure 4.

**Supplement Figure 2:**

Shown are representative medial synovial membrane sections stained with iNOS and NLRP3 in both male and female mice. Quantification is available in figure 4.

**Supplement Figure 3:**Shown are representative knee joint sections stained with p16 (A) and quantification of positive chondrocytes in the (B) medial and (C) lateral side of the tibia. (D) Representative knee joint sections stained with β-Gal and quantification of positive chondrocytes in the (E) medial and (F) lateral side of the tibia in both male and female mice. Males n=19 and females n=19

**Supplement Figure 4:**(A) 3D images of a micro-CT scan of knee joints from male and female mice with different OA severities (cumulative scores at the medial side of the joint). A link to a short movie presenting 3D reconstruction of knee joints with no histological evidence of OA or with high cumulative AC score obtained by histology (Mendeley Data, V1, doi: 10.17632/6nddwstfw3.1)

**Supplement Figure 5:**Femur and tibia SCB traits obtained by micro-CT are significantly directly correlated. Spearman’s rank correlations between SCB BV/TV (A,B), SCB BMD (C,D), SCB Th. (E,F), and SCB Tb.N (G,H) of femur and tibia in male and female mice. Males n=47, Females n=45. P values are indicated for each correlation.

**Supplement Figure 6:**Cumulative AC scores of the medial or lateral tibia show no correlation to the morphology of the SCBP by micro-CT. Spearman’s rank correlations between cumulative AC scores at the medial tibia with SCBP Th (A,B) or SCBP BMD (C,D) in male and female mice. Similar correlations were done between cumulative AC scores at the lateral tibia with SCBP Th (E,F) or SCBP BMD (G,H) in male and female mice. Males n=47, Females n=45. P values are indicated for each correlation.

