## Supplementary figures and images for "Development of primary osteoarthritis during aging in genetically diverse UM-HET3 mice"

### Supplement Figure 1

#### Supplement Figure 1

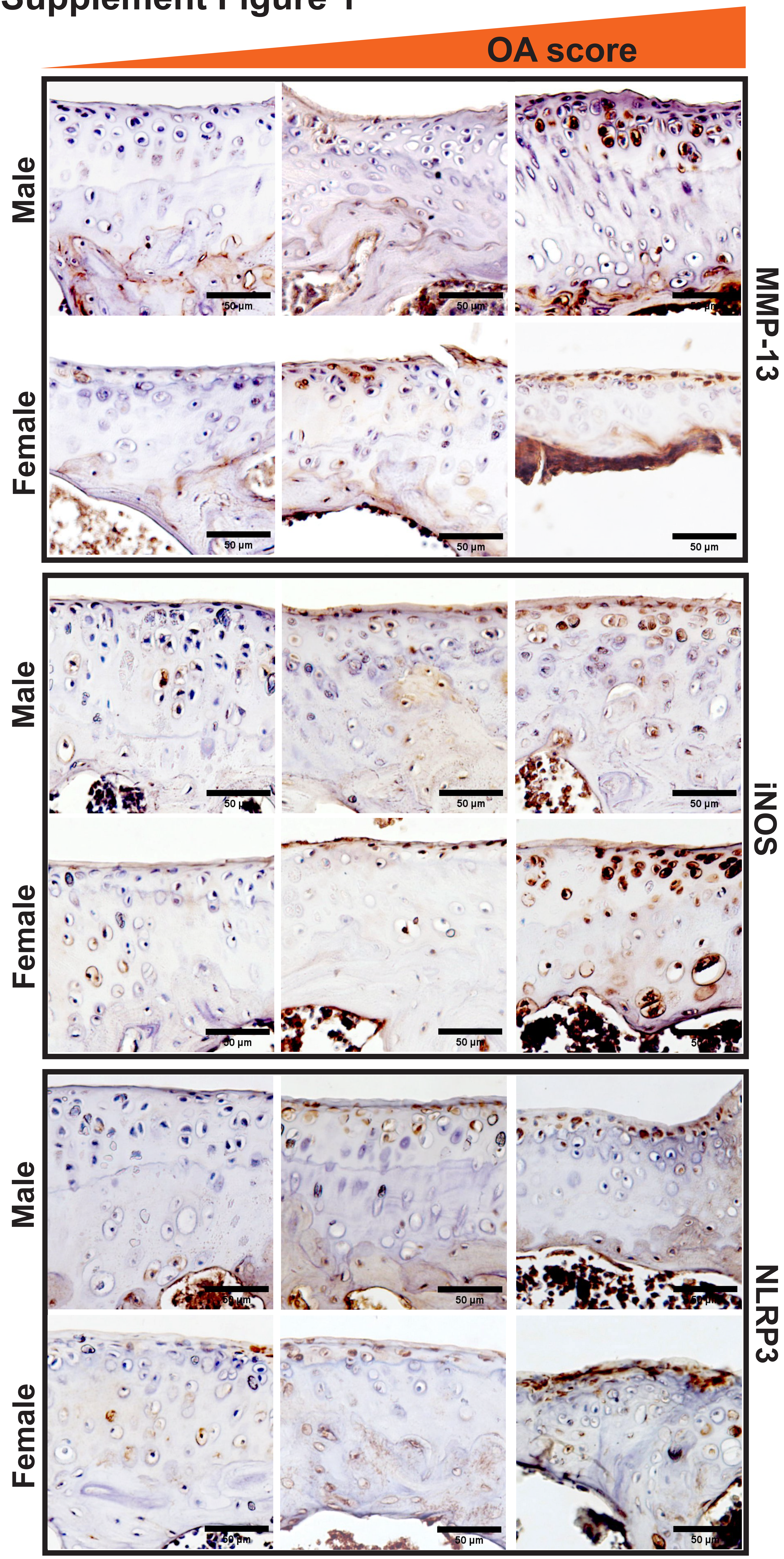

### Supplement Figure 2

#### Supplement Figure 2

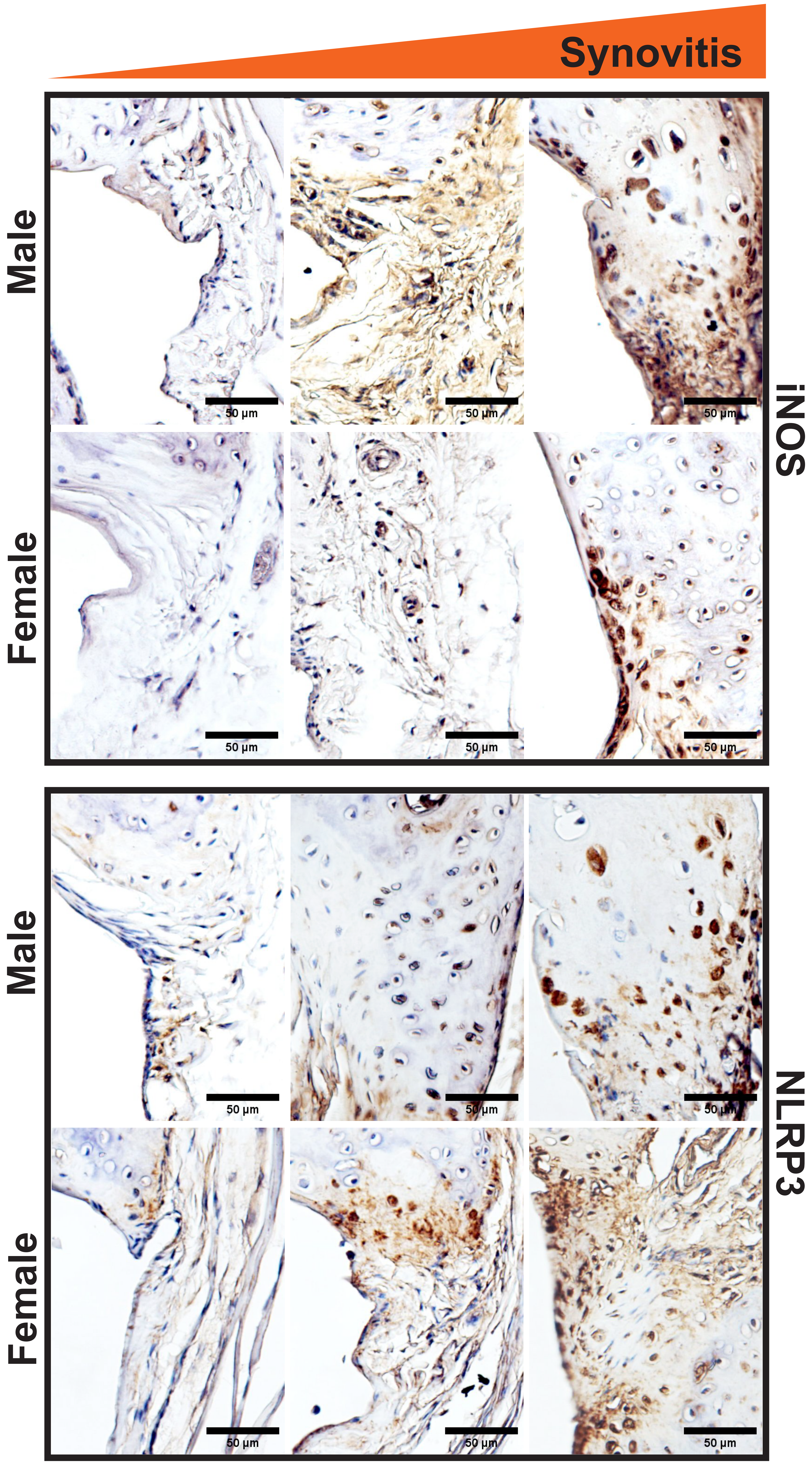

### Supplement Figure 3

#### Supplement Figure 3

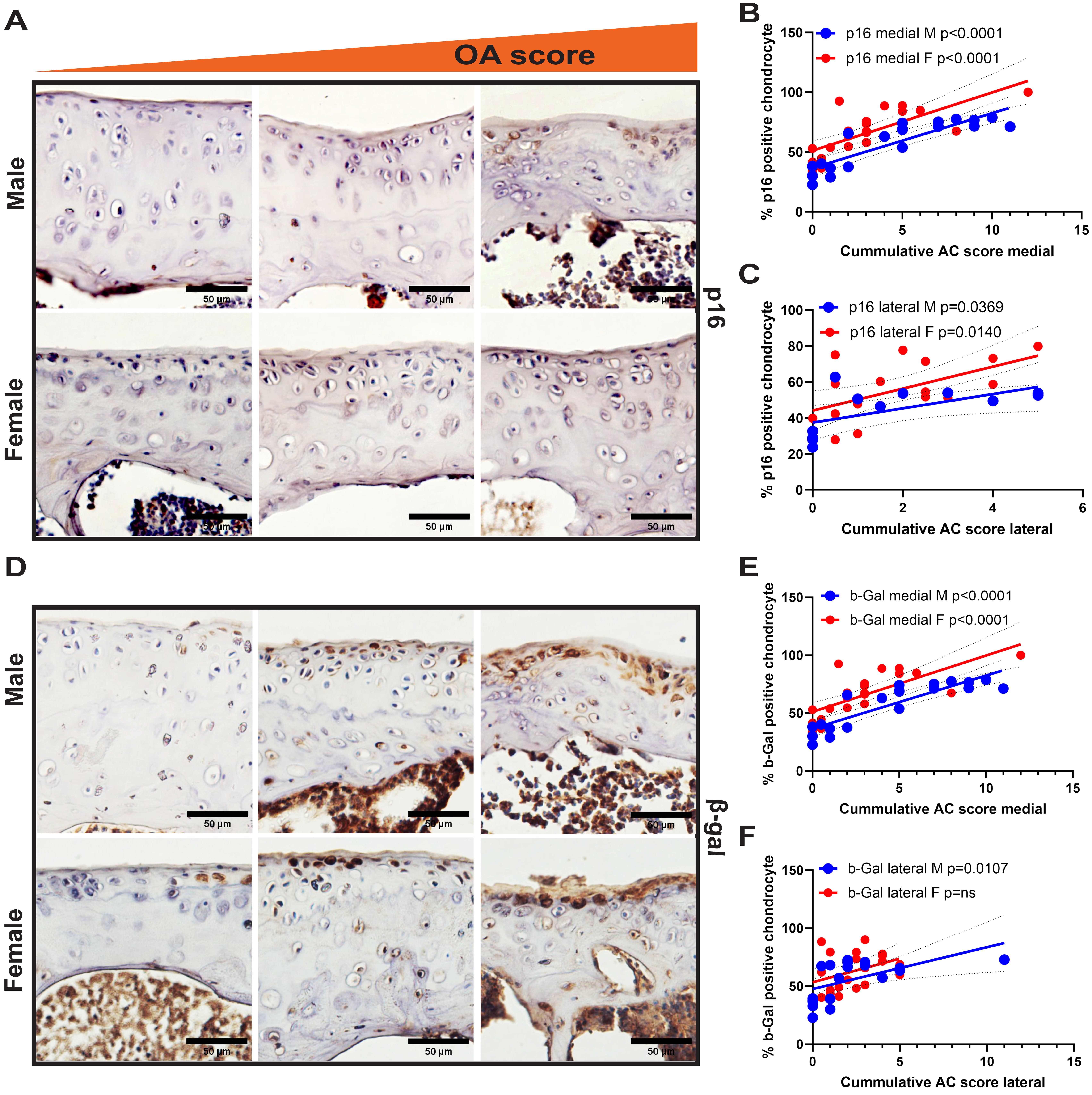

### Supplement Figure 4

#### Supplement Figure 4

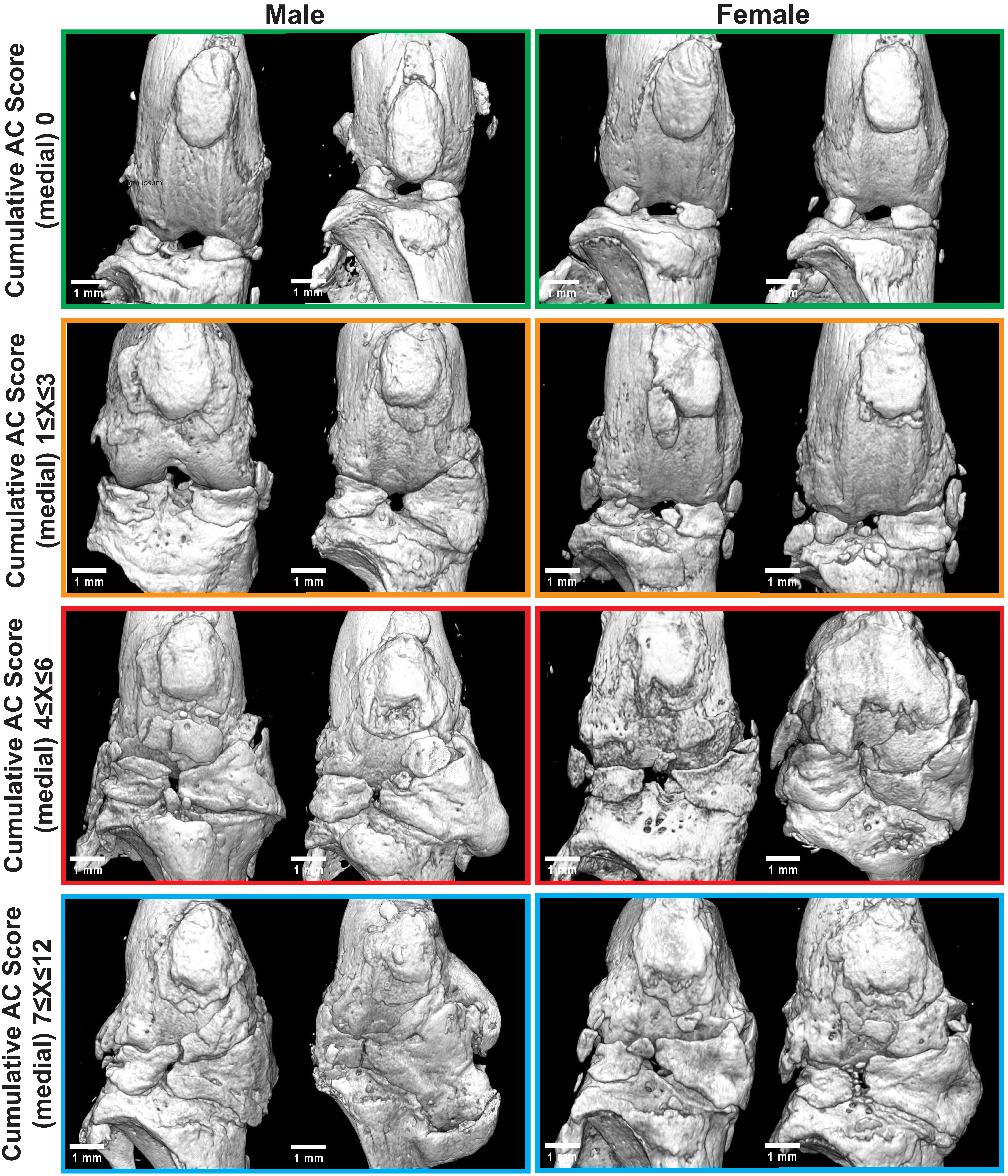

### Supplement Figure 5

Supplement figure 5

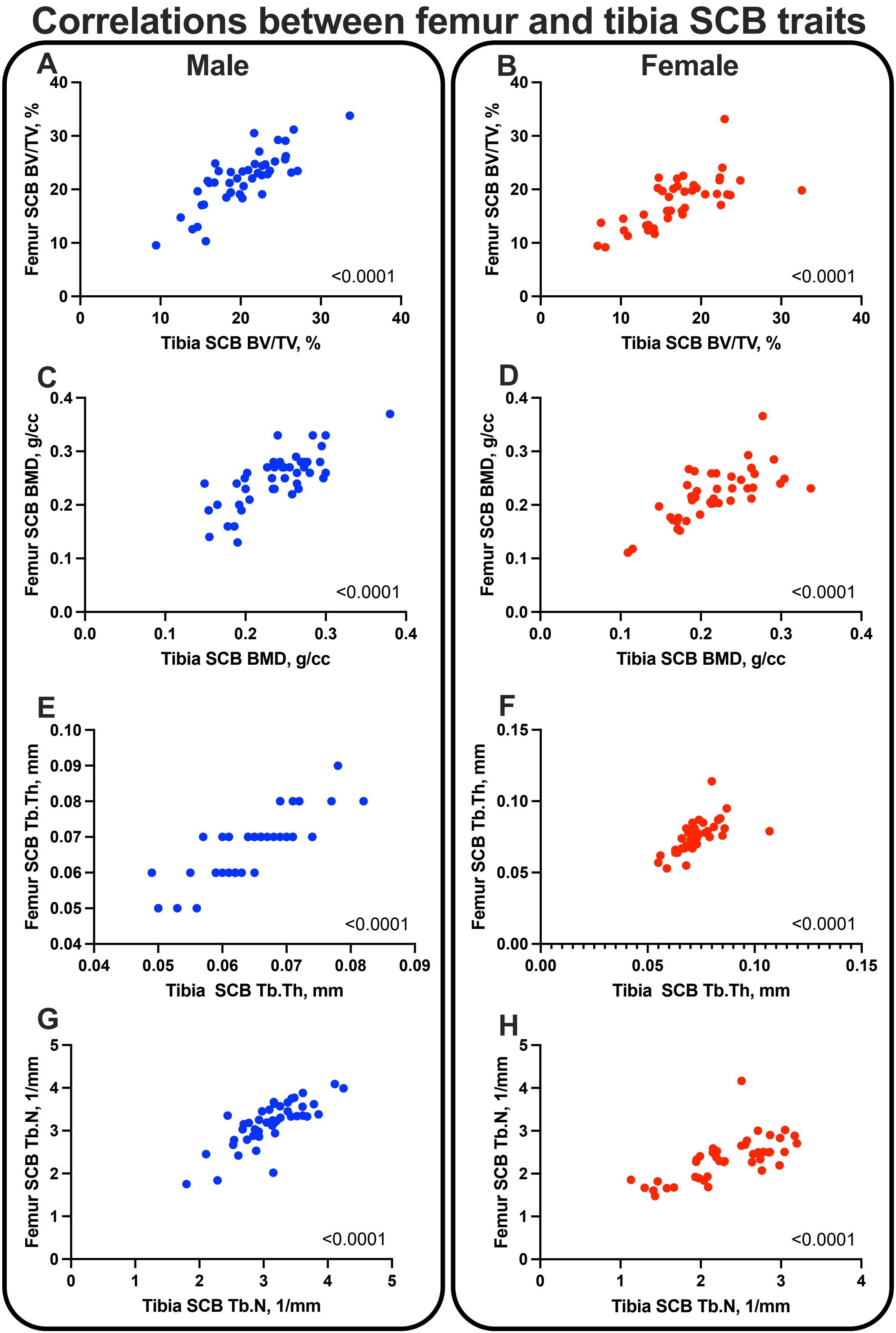

### Supplement Figure 6

Supplement figure 6

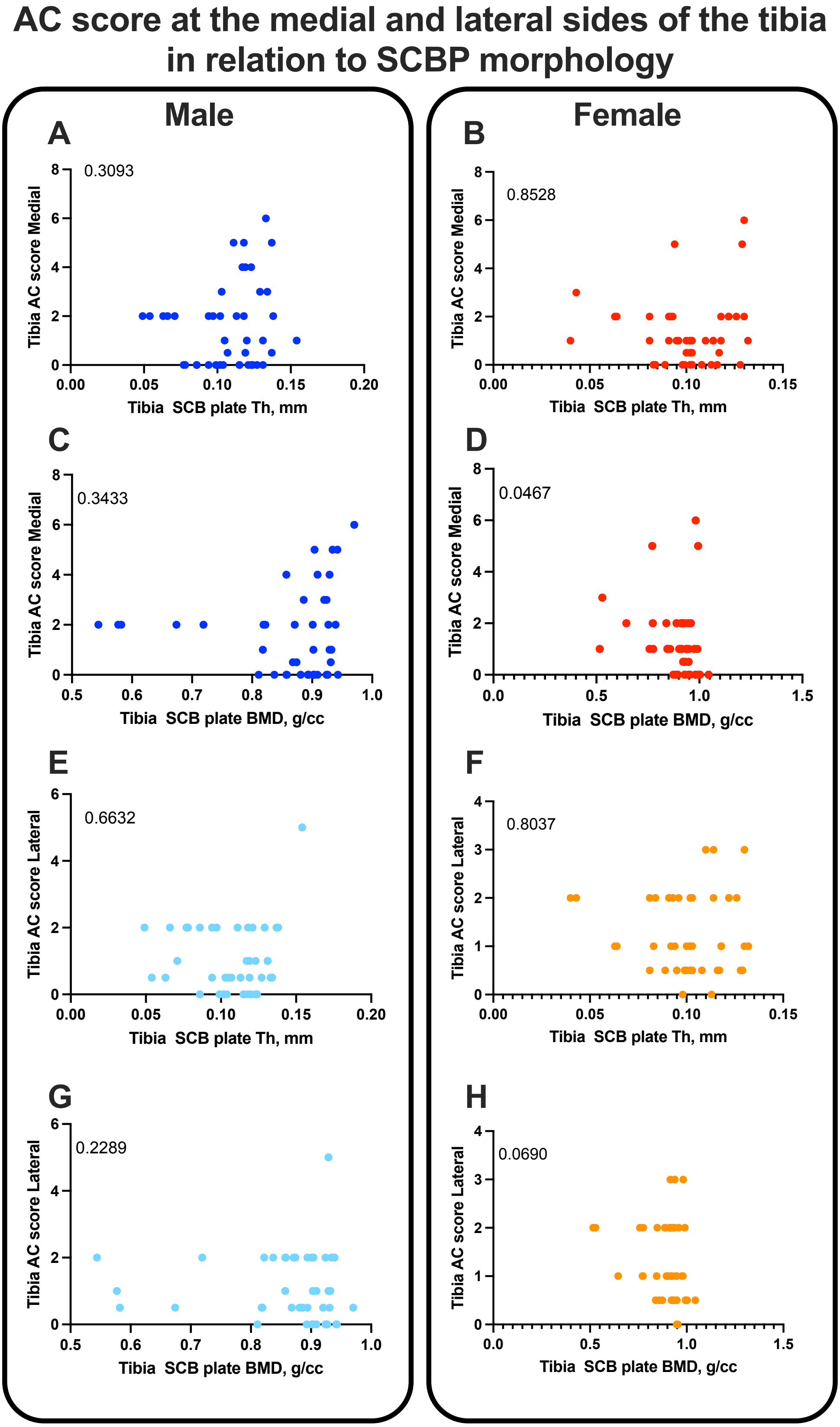
